# TAC-seq: targeted DNA and RNA sequencing for precise biomarker molecule counting

**DOI:** 10.1101/295253

**Authors:** Hindrek Teder, Mariann Koel, Priit Paluoja, Tatjana Jatsenko, Kadri Rekker, Triin Laisk-Podar, Viktorija Kukuškina, Agne Velthut-Meikas, Olga Žilina, Maire Peters, Juha Kere, Andres Salumets, Priit Palta, Kaarel Krjutškov

## Abstract

Targeted next-generation sequencing based biomarker detection methods have become essential for biomedical diagnostics. In addition to their sensitivity and high-throughput capacity, absolute molecule counting based on unique molecular identifier (UMI) has high potential to increase biomarker detection accuracy even further through the reduction of systematic technical biases. Here, we present TAC-seq, a simple and cost-effective targeted <u>a</u>llele <u>c</u>ounting by <u>seq</u>uencing method that uses UMIs to estimate the original molecule counts of different biomarker types like mRNAs, microRNAs and cell-free DNA. We applied TAC-seq in three different applications and compared the results with standard sequencing technologies. RNA samples extracted from human endometrial biopsies were analyzed using previously described 57 mRNA-based receptivity biomarkers and 49 selected microRNAs at different expression levels. Cell-free DNA aneuploidy testing was based on cell line (47,XX,+21) genomic DNA. TAC-seq mRNA biomarker profiling showed identical clustering results to full transcriptome RNA sequencing, and microRNA detection demonstrated significant reduction in amplification bias, allowing to determine minor expression changes between different samples that remained undetermined by standard sequencing. The mimicking experiment for cell-free DNA fetal aneuploidy analysis showed that TAC-seq can be applied to count highly fragmented DNA, detecting significant (*p*=4.8×10^−11^) excess of molecules in case of trisomy 21. Based on three proof-of-principle applications we show that TAC-seq is a highly accurate and universal method for targeted nucleic acid biomarker profiling.

## Introduction

Physiological and pathophysiological disease conditions can be often characterized by the precise quantification of specific nucleic acid biomarkers. There are several methods available enabling detection of RNA- or DNA-based biomarkers but recently, next-generation sequencing (NGS) has become one of the favourite approaches because of its high sensitivity, high-throughput and flexibility. However, despite of its advantages, the relatively high cost of NGS limits its wider application in healthcare. In addition, common NGS assays consist of multiple laboratory steps that often introduce technical biases limiting accurate quantification and therefore hinder the robust and clinically valid detection of biomarkers (Thierry et al. 2014; Wan et al. 2017; Volik et al. 2016). Although quantitative PCR and digital PCR provide simple and cost-effective alternatives to NGS for quantitative biomarker determination, the multiplexing capacity of these approaches is limited compared to highly parallel NGS-based methods.

To overcome the challenges of precise target quantification on extended scale, NGS sensitivity together with high specificity of ligation-PCR have been compiled into common methods and assays. For example, NGS- and ligation-based TempO-Seq (Yeakley et al. 2017) (Templated Oligo assay with Sequencing readout) and MLPA-seq (Kondrashova et al. 2015) are advancement of the well-known MLPA (Schouten et al. 2002) (Multiplex Ligation-dependent Probe Amplification). Both methods overcome original MLPA multiplexing and detection limitations, and enable to apply sensitivity of NGS and analyze up to 20,000 RNA and 200 genomic DNA (gDNA) targets, respectively. Similar approaches are also RASL-seq (Li et al. 2012) for targeted multiplex mRNA analysis and another NGS and targeted ligation-PCR-based method DANSR (digital analysis of selected regions) for cell-free DNA (cfDNA) detection in noninvasive prenatal genetic testing (NIPT). The authors of the latter method report that 384 loci per chromosome 18 and chromosome 21 (altogether 768 loci in a single-tube reaction) are sufficient for aneuploidy discrimination, which makes it significantly less complex assay than so far widely used low-coverage whole genome sequencing for NIPT (Sparks et al. 2012). However, the strategy where ligation-PCR is combined with NGS ensures high level of multiplexing but suffers from the random ligation at low target nucleic acid levels, and on the polymerase-induced errors in both PCR and sequencing steps (Dabney and Meyer 2012; Kebschull and Zador 2015; McInerney et al. 2014). As an outcome, above highlighted methods overestimate the original number of studied molecules and are biased by PCR uneven amplification.

To overcome amplification bias in NGS and to maximize nucleic acid detection sensitivity, <u>u</u>nique <u>m</u>olecular identifiers (UMIs, known also as molecular indexes, unique identifiers or molecular barcodes) are applied. UMI is a string of random nucleotides used in library preparation that was recently introduced to genomic DNA analysis (Kivioja et al. 2011) and also in single-cell transcriptome analysis (Islam et al. 2014). One specific motif out of a large pool of random sequences is incorporated into the original target molecule through oligonucleotides used in library preparation prior to amplification. Later, grouping by identical UMI clones eliminates PCR duplicates and detects the original number of biomolecules. So far, UMIs are widely used in research applications where relatively high PCR amplification is required, such as single-cell analysis and tumor mutation identification (Christensen et al. 2018; Hashimshony et al. 2016; Islam et al. 2014; Kinde et al. 2011; Macosko et al. 2015). Similarly to above-mentioned applications, UMIs can also be used in targeted ligation-PCR NGS assays to enable exact quantification of studied biomarker molecules. Taking the previous into account, we developed a novel ligation- and NGS-based method, TAC-seq, targeted <u>a</u>llele <u>c</u>ounting by <u>seq</u>uencing method for original molecule counting of plasma cfDNA and RNA-based biomarkers.

## Results

### TAC-seq principle and assay design

TAC-seq is a single-tube and ligation-based assay that allows absolute biomarker quantification by the use of two UMI sequences in detector oligonucleotide probes. The studied mRNA and cfDNA molecules are uniquely identified using 54-bp-long target complementary sequence that is detected by two side-by-side located TAC-seq probes (Fig. 1A). Once stringent hybridization of the detector probes to the target occurs, a thermostable ligase is introduced, catalyzing the formation of a phosphodiester bond between the 5’-phosphate and the 3’-hydroxyl of two side-by-side detector probes. Next, ligated detector-target complexes are captured using magnetic beads and amplified by PCR (Supplemental Fig. S1), resulting in a ready-to-sequence library within a 3 h turnaround time. The risk of losing alleles is minimized by the dilution-free protocol in which ligated detector probes are captured, amplified and identified by sequencing. To simplify the *in silico* design of specific TAC-seq probes and data analysis, we developed an online tool for designing the TAC-seq probes (http://nipt.ut.ee/design/) and a computational workflow software to enable data processing (open-source software link is in Methods and principle shown in Supplemental Fig. S2).

**Figure 1.**
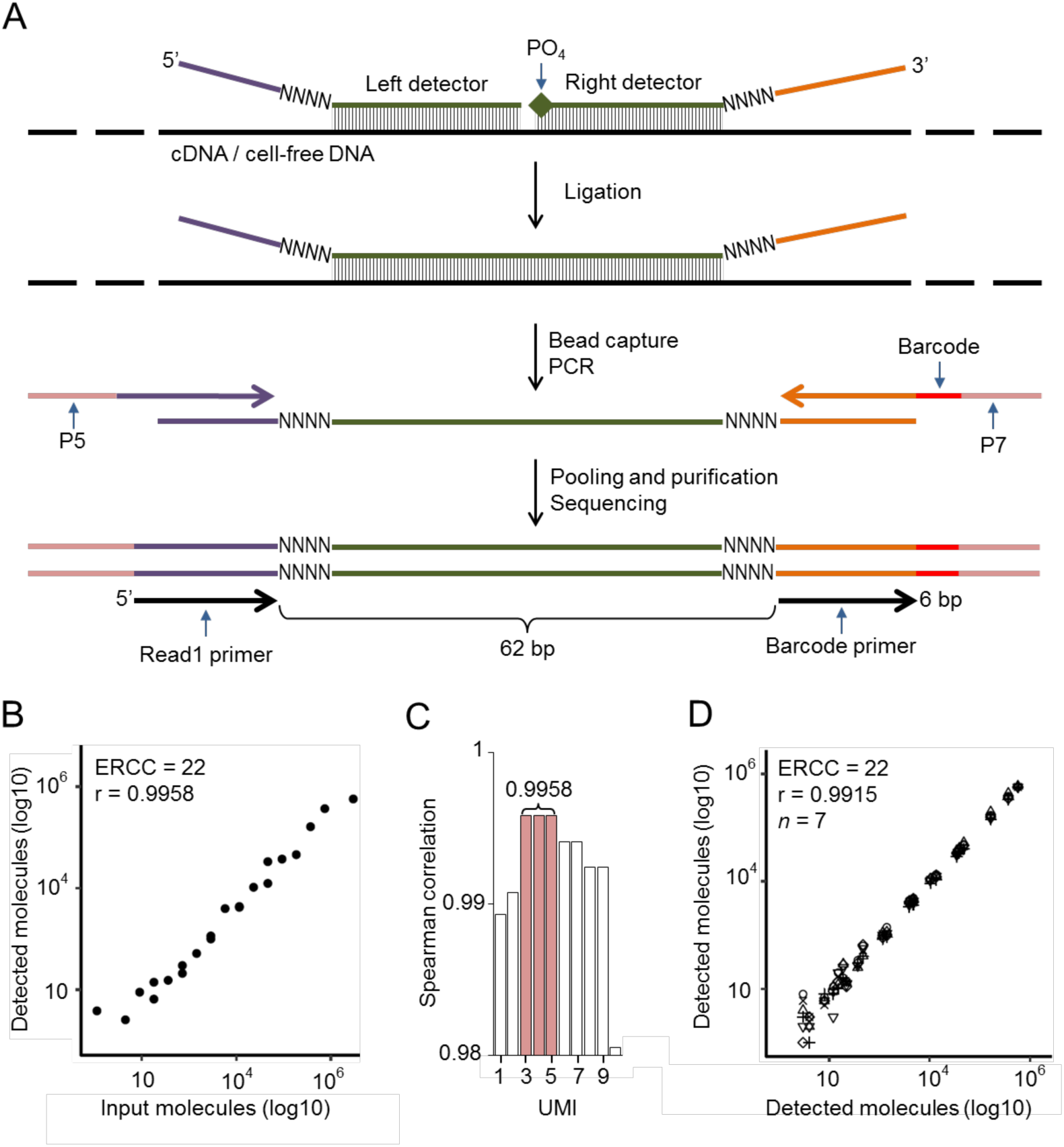
Principle and technical parameters of TAC-seq. (*A*) Schematic diagram of the assay to detect specific mRNA or cell-free DNA. Target-specific DNA oligonucleotide detector probes hybridize under stringent conditions to the studied cDNA or cfDNA. Both detector oligonucleotides consist of a specific 27-bp region (green), 4-bp unique molecular identifier (UMI) motif (NNNN) and universal sequences (purple and orange). The right detector oligonucleotide is 5’ phosphorylated. After rigorous hybridization, the pair of detector probes is ligated using a thermostable ligase under stringent conditions. Next, the ligated detectors complexed with the target region are captured with magnetic beads and PCR amplified to introduce sample-specific barcodes and other common motifs that are required for single-read NGS. (*B*) Spearman correlation analysis of the input and detected ERCC synthetic spike-in mRNA molecules at UMI threshold 4 (UMI=4). UMI threshold is defined as the number of detected unique UMI sequences. For example, UMI=4 indicates that a certain UMI motif is detected at least four times. UMIs are valuable only if the number of UMI combinations (8-bp UMI provides 65,536 variants, for example) is substantially larger than the sum of the target molecules in the studied sample. (*C*) Bar plot of Spearman’s correlation analysis of the ERCC input and detected molecules at different UMI thresholds. (*D*) Reproducibility of seven technical ERCC replicates (seven different icons on plot) of 22 spike-in molecules at UMI=4.

### Experimental evaluation of TAC-seq

First, we used External RNA Controls Consortium (ERCC) RNA spike-in controls to validate the technical sensitivity and accuracy of the TAC-seq method. Altogether, 22 spike-in sequences were assayed at various concentrations, ranging from 1 to 3×10^5^ molecules per reaction (Supplemental Table S1). The spike-in sequences were then detected with TAC-seq probes, sequenced and counted at different UMI thresholds. The analysis consistently demonstrated a high correlation (Spearman *r*>0.99, Fig. 1B–C) between input and detected molecules for both relaxed and conservative (*n*>1 molecules required) UMI thresholds (Shugay et al. 2017). These results suggested that conservative UMI thresholds (*n*≥3 molecules required in this case, Fig. 1C) are justified and applicable for high-coverage sequencing, in which the unfiltered read numbers are significantly higher than the UMI corrected outcome (Shugay et al. 2017). With seven technical ERCC replicates, the average 1.5×10^6^ raw read count per replicate dropped to 5.7×10^3^ after UMI correction, demonstrating a 102-fold average PCR redundancy at UMI threshold of 4 (*n*=4) (Supplemental Table S1). Additionally, excellent reproducibility (Spearman *r*=0.9915, Fig. 1D) among seven ERCC replicates was demonstrated.

### mRNA detection – absolute molecule counting of endometrial receptivity biomarkers

Next, we designed a transcriptome assay to analyze human endometrial linings. We targeted 57 endometrial receptivity mRNA transcripts that are potential biomarkers in reproductive medicine for testing embryo implantation compatibility (Altmae et al. 2017). Ten endometrial biopsy samples were analyzed by TAC-seq and compared with the levels of selected 57 transcripts from full transcriptome RNA sequencing (RNA-seq) data. Principal component analysis of these data showed identical clustering of samples when applied to RNA-seq results and two different TAC-seq assays, carried out with high-(denoted as TAC-seq_high_, average 25.7×10^6^ reads per sample) and low-coverage (TAC-seq_low_, average 1.23×10^6^ reads per sample) sequencing (Fig 2A). In these analyses, the first component described most of the sample variability (RNA-seq 89.8%, TAC-seq_high_ 79.6%, TAC-seq_low_ 83.0%) and clearly distinguished the pre-receptive and receptive samples (*n*=5 in both groups, Fig. 2A), except for one outlier sample both in RNA-seq and TAC-seq analyses. The same sample grouping was confirmed by hierarchical clustering: four pre-receptive samples clustered together with high confidence (approximately unbiased (AU) probability of 100%), and receptive samples clustered together with one outlier sample in all datasets (*AU*_*RNA*_=94%; *AU_TAC-high_*=69%; *AU_TAC-low_*=81%) (Fig. 2B and Supplemental Fig. S3).

**Figure 2.**
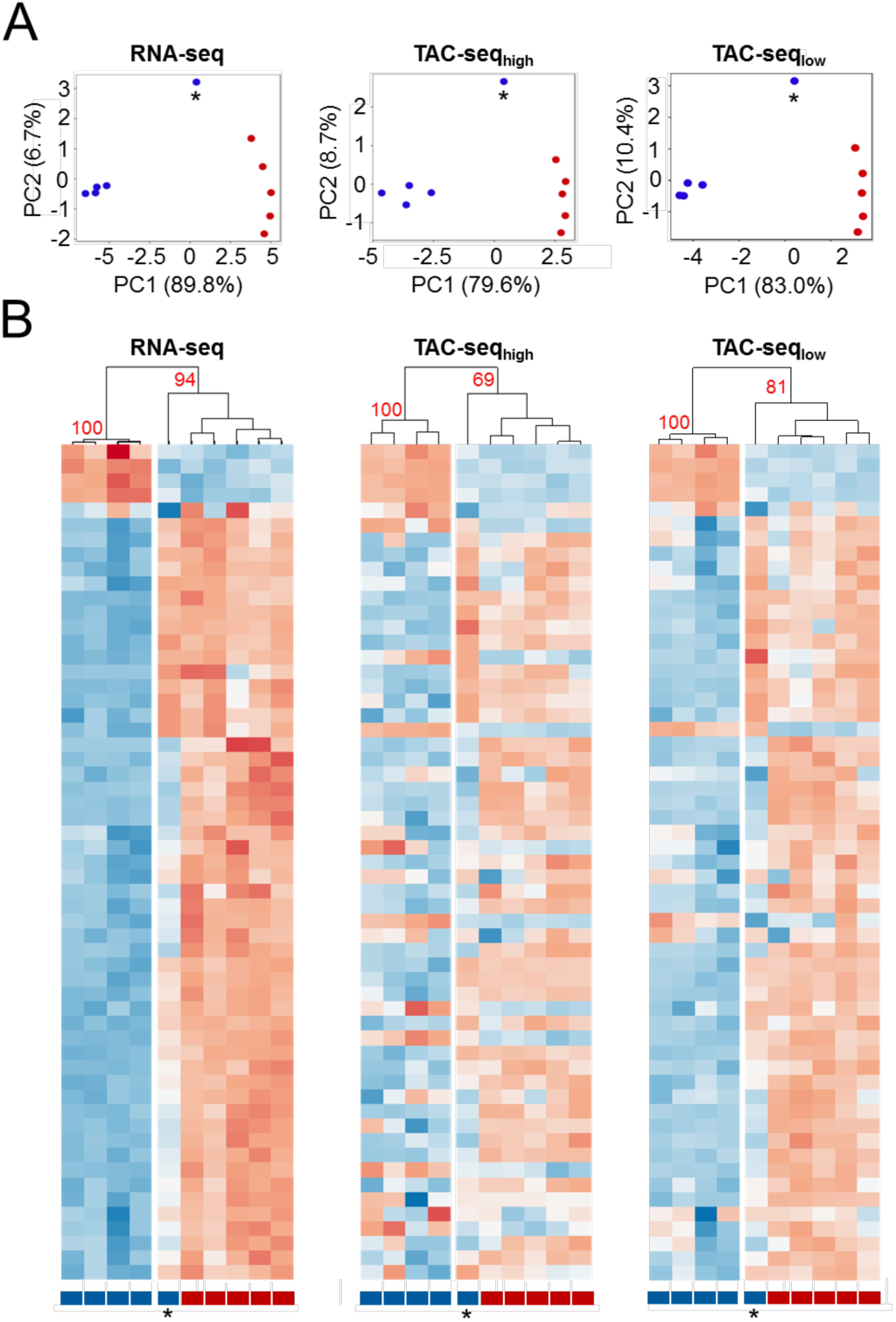
Comparison of the overall predictions for mRNA TAC-seq assay. (*A*) Principal component analysis of the full transcriptome RNA-seq, high-coverage TAC-seq and low-coverage TAC-seq of 10 endometrial samples. The first principal component (PC1) describes most of the sample variability and correlates most with the receptivity status. Blue dots represent pre-receptive and red dots receptive human endometrial samples. One separate pre-receptive sample (indicated with an asterisk) represents the same sample that clusters differently in the heatmap analysis (below) and is therefore a potential biological outlier. (*B*) Heatmaps of the full transcriptome RNA-seq, high-coverage- and low-coverage TAC-seq show the sensitivity to distinguish different endometrial samples according to their receptivity. One pre-receptive sample (indicated with an asterisk) shares the expression profile and clusters together with receptive samples in all three comparisons. Pre-receptive samples are labelled blue and receptive red. Detailed heatmaps are presented in Supplemental Fig. S3 together with housekeeping genes that demonstrate a lack of fluctuation of the pre-receptive and receptive biopsies. High-coverage TAC-seq data are presented at UMI=2 and low-coverage data at UMI=1 on PCA and heatmaps. The data are plotted as row-wise scaled log-transformed counts per million (CPM) values. The samples are hierarchically clustered column-wise using Pearson correlation. The genes are ordered row-wise according to the RNA-seq clustering results using Euclidean distance. Fewer genes are found expressed with a low-coverage compared to RNA-seq and high-coverage TAC-seq.

In both high- and low-coverage TAC-seq approaches all 57 selected biomarkers were detected, demonstrating a flawless detection across all differentially expressed transcripts. However, high-coverage sequencing analysis revealed that 8-bp UMI causes technical limitation in molecule detection in the upper detection range. Six highly expressed genes met UMI-related saturation and were not precisely quantified (Supplemental Fig. S4). This technical limitation can be easily overcome by adding longer UMI nucleotides to detector oligonucleotides or by sequencing with lower coverage.

In low-coverage assay we expanded the assay to a 70-plex by adding housekeeping genes (8) and selected ERCC spike-ins (5). We observed that although the assayed housekeeping genes represented 47.4% of all unique reads in this assay, the biomarker-based clustering probability was still high (*AU_TAC-low_*=81%) and all of the targeted 57 biomarkers were detected (Fig. 2B). Based on these results, we suggest that highly expressed housekeeping genes (eg. *ACTB, GAPDH* and *PPIA*) could be excluded from the low-coverage assay, to ensure accurate transcriptome biomarker profiling with only 500,000 sequencing reads.

### microRNA detection – differentially expressed molecules in endometrium

To test the feasibility of TAC-seq with small non-coding microRNAs (miRNA), we selected 49 miRNAs from endometrial tissue, which were previously analyzed by small RNA-seq and covered highly variable expression levels from 10 to 4,712 counts per million. TAC-seq detected all 49 assayed miRNAs over 16 analyzed endometrial samples with high reproducibility (Spearman *r*>0.997, Fig. 3) and showed high sensitivity in order to distinguish biologically different pre-receptive and receptive clinical biopsies in unsupervised clustering that were not detected by previous RNA-seq assay (Supplemental Fig. S5).

**Figure 3.**
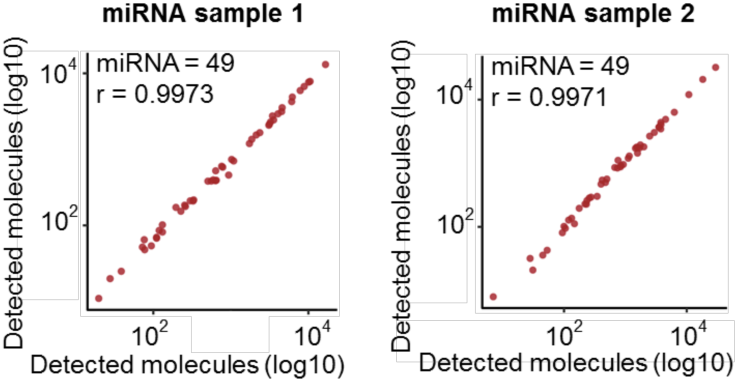
TAC-seq miRNA assay performance. Correlation plots of four miRNA sample technical replicates using TAC-seq assay at UMI=4. miRNA sample 1 is on the left hand and has two replicates, one plotted on the X-axis and the other on the Y-axis. The same with miRNA sample 2 on the right hand.

Due to the nature of miRNAs, short, 20–24-bp target regions were detected using specific probes, and the TAC-seq protocol was modified accordingly (Supplemental Fig. S6). Additionally, an *in silico* prediction based on RNA-seq suggested that the selected set of miRNAs had an extreme nucleotide imbalance (Starega-Roslan et al. 2015) at positions one (G<2%) and five (C<3%), causing potential sequencing failure (Supplemental Fig. S7). The limitation was overcome using an adjusted spike-in probe that was designed to compensate for the two less-represented nucleotides at certain positions and therefore balance the whole sequencing run. The custom spike-in was later added to the sequencing reaction, resulting in a balanced nucleotide distribution and a high, 96% pass-filtering read rate (Supplemental Fig. S8). Although RNA-seq is convenient for miRNA profiling, our results confirm the previous findings (Faridani et al. 2016) that amplification bias reduction using UMI can enhance sensitivity. As a result, TAC-seq allowed clear separation of biologically different samples even in case of minor expression differences (Supplemental Fig. S5).

### Cell-free DNA detection – trisomy detection using controlled conditions

As the last test, we evaluated the potential of the TAC-seq method for NIPT, a widely used NGS-based clinical application. NIPT detects fetal trisomy based on the molecular counting of fetal cfDNA in maternal blood samples. We designed a proof-of-principle trisomy 21 detection assay to demonstrate the applicability of the TAC-seq method for absolute cfDNA molecule counting to determine fetal trisomy at different experimentally controlled ‘fetal fraction’ levels. In the following *in vitro* aneuploidy detection experiment, we mixed different proportions of gDNA from a chr21 trisomy cell line with a known normal control gDNA. Different proportions of fetal trisomic cfDNA were created to imitate the range of fetal fraction in mothers’ circulating cfDNA. gDNAs were sheared by sonication to mimic 160–180 bp cfDNA, mixed to yield 5–30% trisomy proportions and hybridized with TAC-seq detector probes that were designed to target 114 loci along chr2 and chr21. Normal diploid chr2 served as a reference. The molecule counts without outliers detected a significant (*p*=4.8×10^−11^, one-tailed Welch’s t-test) excess of aneuploid chr21 reads compared to normal chr2 in both trisomy 21 cell line replicates (Fig. 4A).

**Figure 4.**
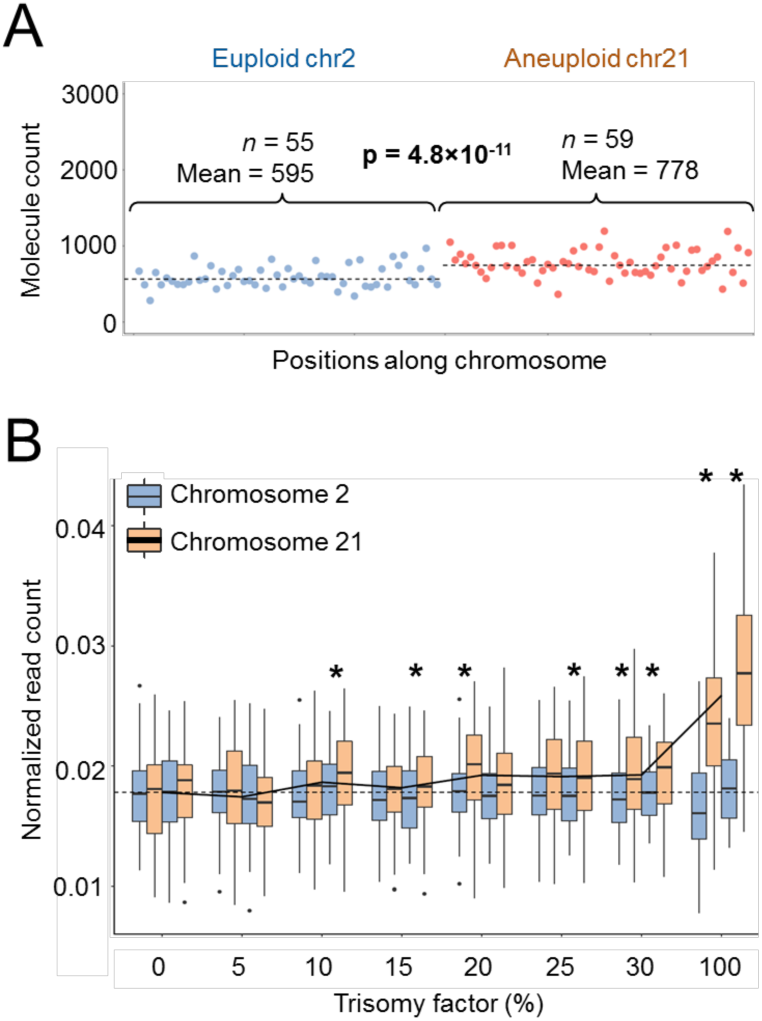
Trisomy detection. (*A*) Selected 114 loci along chr2 and chr21 based on their absolute molecule counts. Scatter plot for euploid chr2 (blue) and aneuploid chr21 (orange) DNAs with the mean values clearly distinguishing (p = 4.8×1 O’^11^, one-tailed Welch’s t-test, two replicates) the trisomy 21 cell line’s DNA with excess detected molecule count at UMI=1. (ß) Boxplot of normalized molecule counts of *in vitro* trisomy TAC-seq indicates a positive correlation between the trisomy factor (trisomie cell proportion) and chr21 counts. There were two biological replicates for each trisomy factor plotted side-by-side. The dashed line represents the mean value of normalized molecule counts of chr2 for normal reference samples (trisomy factor = 0%). The solid black line represents the mean value of normalized molecule counts of chr21 for different trisomy factor percentages. The asterisks indicate significant chr2 and chr21 read-count based differences between studied samples (p<0.05, one-tailed Welch’s t-test).

The detection of trisomy 21 demonstrated a significant difference at the lowest 10% ‘fetal cfDNA fraction’ in one replicate out of two, which improved even further in case of every additional 5% increase in the ‘fetal fraction’. At the 30% ‘fetal fraction’ level and with a pure chr21 trisomy cell line, both replicates detected a significant difference (Fig. 4B). The same result was confirmed by an independent whole-genome re-sequencing NIPT assay for the same samples (Bayindir et al. 2015) (Supplemental Fig. S9). Interestingly, we observed that non-normalized molecule counts showed different median values for chr2 and chr21 in a normal diploid cell line, with the chr2 molecule counts being, on average, 5.3% higher than those of chr21. The trend was confirmed with two independent samples.

## Discussion

Combining high sensitivity and flexibility of NGS with cost-efficient and precise quantification of targeted methods can enable robust detection of specific nucleic acid biomarkers indicative of (patho)physiological conditions. TAC-seq is an advanced ligation-based NGS method that differs from existing ligation-PCR assays such as MLPA (Schouten et al. 2002), MLPA-seq (Kondrashova et al. 2015), TempO-Seq (Yeakley et al. 2017), RASL-seq (Li et al. 2012) and DANSR (Sparks et al. 2012). The major advantage of TAC-seq is ability to detect the number of original molecules of transcriptomic biomarkers such as mRNA and miRNA, and genomic loci from cfDNA. Precise molecule counting is achieved by integrated UMI or ‘molecular barcode’ motif (Kivioja et al. 2011) which decreases the quantitative and random bias introduced by *in vitro* replication steps. Using UMIs removes PCR duplicates, reducing one of the major NGS-specific technical biases and improves the accuracy of NGS.

We detected very high sensitivity correlation over 22 analyzed ERCC spike-in input and detected molecules (Spearman *r*=0.9958 on Fig. 1D) with high coverage, ensuring that each UMI coverage was 102×. Based on the coverage, we are confident that very few UMIs have been missed and therefore this outcome is reliable. However, systematic difference appears between lowly- and highly-expressed targets with the number of high copy-number molecules is underestimated (see top-four ERCC spike-ins in Supplemental Table S1). This is explained by the length of UMI sequences, causing ‘technical saturation’. Eight nucleotide UMIs used in our study have 65 thousand possible sequences, which is well suitable for cfDNA-based trisomy detection as copy numbers of cfDNA in 10 ml of blood remains <5,000 (Kukita et al. 2015; Newman et al. 2016). The same applies to TAC-seq expression applications if lower RNA concentrations are used. Alternatively, it is possible to extend UMI sequences in both detector probes from current 8 to 12 nucleotides, ensuring 16.7 million possible combinations. At the same time we are aware that introducing significantly longer and random UMI strings into detector probes may increase the probability of probes self-pairing and unspecific ligation. However, UMI-related issues such as ‘saturation’ and replication-caused novel, ‘phantom’ UMIs (Smith et al. 2017) that should be taken into account in assay design and data analysis.

TAC-seq was designed while keeping in mind the main prerequisites of genetic testing laboratories – sensitivity, robustness and cost-efficiency. Sensitivity and molecule counting by UMI was discussed above. Robustness is ensured by single-tube protocol to minimize the risk of allelic drop-out. Furthermore, the approach is dilution-free, meaning that analyzed biomarker molecules together with ligated detector probes are captured and identified by sequencing. The latter is crucial in liquid biopsy samples where each locus is represented only by some thousands of copies. If ligation-based assay with specific probes is used then probe hybridization-compatible target cfDNA copy-number reduces 25% due to the short length of cfDNA (180 bp) as the locus is not detected if it locates closer than 25 bp to cfDNA fragmentation site.

TAC-seq detects mRNA biomarkers through oligo-T primed cDNA synthesis (poly-A selection) that reflects the analysis of active transcriptome. It differs from TempO-Seq (Yeakley et al. 2017) where recently described SplintR ligase (Lohman et al. 2014) for RNA/DNA hybrid is used to detect any, even fragmented RNA targets by specific detector oligonucleotides. In addition, SplintR ligase optimum working temperature is up to 37˚C that may limit the specificity of formed RNA/DNA probe complexes prior ligation. In contrast, TAC-seq uses thermostable *Thermus aquaticus* (*Taq*) DNA ligase (Housby et al. 2000) that enables highly specific hybridization and ligation at temperature over 45˚C. Based on this property of *Taq* DNA ligase, we have carried out first specific probe-target hybridization at 60˚C and thereafter introduced ligase to join the proximity annealed strands at the same temperature.

As sequencing contributes to the majority of the NGS cost, it is critical to apply library preparation that supports low-coverage sequencing in routine NGS clinical applications. Cost-effectiveness of TAC-seq is ensured by off-the-shelf reagents and the usage of common instruments in genomic laboratory, such as standard thermocycler and benchtop NGS sequencer. The running cost of TAC-seq is only a fraction of the cost for commonly used NGS applications like whole-genome sequencing for NIPT or RNA-seq for mRNA and miRNA analysis. Setup cost of TAC-seq depends on number of studied loci due to the need of specific detector oligonucleotides (Supplemental Fig. S10). Consumables and their approximate prices are listed in Supplemental Table S2 and explained in Methods. Based on our in-house library preparation and sequencing, the total reagent cost for miRNA profiling and cell-free DNA analysis is less than 30 EUR per sample and 26–40 EUR per sample for mRNA biomarker analysis, depending on the sequencing depth. Therefore, TAC-seq has the potential to become a cost-effective alternative for routine NIPT after clinical studies or for detecting the levels of transcriptome biomarkers.

TAC-seq probe specificity is ensured by 54 bp-long region on mRNA and gDNA. We developed automated mRNA probe design software (http://nipt.ut.ee/design/ without

restrictions in usage and described in Methods) that automates probe design procedure and provides highly specific oligonucleotide sequences with common motifs that are ready for synthesize. Probe design for miRNA molecules is even more straightforward and does not require special software (see Supplemental Fig. S6). Another simplification that makes NGS as a choice of detection method is user-friendly data analysis. Small-scale NGS data analysis does not demand powerful computing resources. For this, we provide user-friendly personal computer software for small-scale TAC-seq data analysis and open source code for intense analysis (link in Methods). The NGS ‘big data’ limitation has been overcome by simple analysis pipeline. Most resource-demanding raw data processing after sequencing is done by Illumina^®^ cloud-computing environment. Following TAC-seq analysis is based on text-file manipulations eliminating the need of sequencing read mapping, making it possible to perform NGS analysis in personal computer (see details on Methods).

In conclusion, we have developed a novel and highly parallel method to count accurately the number of nucleic acid biomarker molecules in studied samples. Our proof-of-principle study demonstrates that TAC-seq has similar sensitivity to golden standard RNA-seq method in case of mRNA and miRNA application, and can successfully detect the excess of cfDNA molecules (indicative of chromosomal trisomy) in cfDNA-like material. TAC-seq is automation-compatible method that is designed to overcome ligation- and NGS-based limitations in genetic testing laboratory. Although all presented applications need careful clinical validations before they can be utilized, the described method is the base for further specialization and optimization to provide advanced DNA and RNA biomarker screening tools and thereby improve reach and quality of corresponding research and healthcare applications.

## Methods

### Studied samples

This study was approved by the Research Ethics Committee of the University of Tartu (246/T-21 and 221/M-31). Endometrial biopsies for RNA assays were collected from healthy volunteers with proven fertility as previously described (Altmae et al. 2017). Genomic DNAs from GM01359 (47,XY,+18) and GM04616 (47,XX,+21) cell lines (NIGMS Human Genetic Cell Repository, Coriell Institute of Medical Research) were extracted using the DNeasy Blood and Tissue kit (Qiagen). To mimic cfDNA, genomic DNA was fragmented to 150-200 bp using Covaris M220 Focused-ultrasonicator (Thermo Fisher) with the following settings: 50 µl volume, 10% duty cycle, 75 W peak incident power, 200 cycles per burst and 360 s. DNA concentrations were quantified using Qubit dsDNA HS (Thermo Fisher).

### Biomarker selection and TAC-seq probe design

The biomarkers of endometrial receptivity were selected based on our previous publication (Altmae et al. 2017). Briefly, nine studies including a total of 164 endometrial samples from fertile women were included in a meta-analysis using the Robust Rank Aggregation method. In the current study, we used the 57 mRNAs identified as potential endometrial receptivity biomarkers for distinguishing the pre-receptive and receptive endometrial samples. A pair of TAC-seq detector DNA oligonucleotide probes (left detector and right detector) was designed for every targeted gene using the special TAC-seq probe design software (http://nipt.ut.ee/design/). All of the oligonucleotides used in this study are listed in Supplemental Table S3. Both the left and right detectors consisted of a specific sequence (27-bp), an UMI (4-bp) and a left universal sequence or right universal sequence. Each detector pair targeted the coding sequence in the Consensus Coding Sequence Set (CCDS). For genes without CCDS, the most likely transcript was chosen manually from the Ensembl 87 database. Selection of the target sequence was based on two criteria. First, the adjacent 14-bp, 7-bp from both detector probes around the ligation site, had to be unique against human cDNA (GRCh38) to minimize the likelihood of non-specific hybridization. Next, the unique sequences were ranked according to the distance from the 3’-end of the transcript. Routine genetic testing detectors were preferentially designed close to transcript’s 3’-end to minimize the effect of possible RNA degradation caused by sampling and handling if poly-A at the mRNA 3’-end is used for cDNA priming. Additionally, detector-specific regions were filtered by GC-content to determine the optimal melting temperature. The overall GC-content of a probe had to be between 40–60%, and the GC-content of the adjacent ends (4-bp) was up to 50%. Additionally, detector oligonucleotides with inter- or intra-complementarity issues were excluded from the selection. Although mRNA’s 3’-ends were targeted in this study, the software has an option to design TAC-seq detectors close to the transcript’s 5’-end, if required. The ERCC spike-in 22 detectors were designed based on the above description close to their poly-A tails.

For the TAC-seq miRNA assay, 49 miRNAs showing stable expression values (standard deviation/mean count per million (CPM) <0.5) within a study group were chosen according to previously published small RNA sequencing data (Altmae et al. 2017). One specific 20–24-bp detector oligonucleotide was designed per each selected miRNA (‘Specific detector’ in Supplemental Fig. S6). Eight UMI nucleotides and a common sequence were added to each specific detector probe. The right detector oligonucleotide is universal for all miRNAs, consisting of two common sequences and a 5’ phosphate to enable ligation.

Chromosome 2 and 21 loci were selected from the k-mer http://bioinfo.ut.ee/NIPTMer/programs/lists/ database (converted to text files with glistquery http://bioinfo.ut.ee/NIPTMer/programs/glistquery where k-mers overlapping known polymorphisms (dbSNP build ID 149) were first removed and the remaining candidates were used as an input for BLAST 2.4.0+ (task blastn) with database version GRCh38 (GCA_000001405.15). All reads with more than one exact match were removed, following the concatenation of overlapping regions. The regions were converted to sequences with UCSC Genome Browser Gateway. Altogether, 114 specific detector pairs over the studied chromosomes 2 and 21 were selected according to the above-described design, ensuring equal coverage over the entire chromosome.

### ERCC mRNA library preparation

Non-skirted low profile PCR Strip Tube Plates (Thermo Fisher) were used with domed cap strips (Thermo Fisher). ERCC Spike-In Mix 1 (Life Technologies) was first diluted 10× and then additionally 100× with water. Aliquots, each containing 1.3 µl of 1,000× dilution, were stored at −70°C until use. Next, 199 µl of water was added to 1.3 µl aliquot and mixed. 1 µl of diluted ERCC spike-in content (Supplemental Table S1), serving as a template for each individual library was added to 2 µl of denaturation buffer, containing 5 mM Tris-HCl (pH 7.0) (Sigma-Aldrich), 1 mM dNTP mixture (Thermo Fisher), 400 nM Oligo-T30 primer and 0.05% Triton X-100 (Sigma-Aldrich). Reaction was mixed by pipetting and centrifuged briefly. RNA was denatured by 1 min at 80°C and immediately placed on ice. After that, reverse transcriptase (RT) master mix containing 100 mM Tris-HCl (pH 8.5) (Sigma-Aldrich), 2.5 M betaine (Sigma-Aldrich), 150 mM KCl (Sigma-Aldrich), 10 mM DTT (Sigma-Aldrich), 15 mM MgCl_2_ (Sigma-Aldrich), 4 U RiboLock RNase inhibitor (Thermo Fisher) and 20 U Maxima H Minus Reverse Transcriptase (Thermo Fisher) was prepared. The master mix was vortexed and briefly centrifuged. 2 µl of RT master mix was added to previously denatured RNA (3 µl). All RT pipetting steps were performed on ice. cDNA synthesis was performed by 30 min at 42°C, following 5 min at 85°C for RT inactivation.

Twenty two TAC-seq detector pairs targeting ERCC spike-in molecules were previously mixed together from 100 µM stock solutions, creating a ‘100 µM’ oligo pool. The oligo mixture was diluted to 5 µM by water and stored at −20°C. Once cDNA synthesis was completed, 1 µl of 5 µM TAC-seq detector mixture was added to RT mixture. The content was mixed on vortex and centrifuged briefly. Strip tubes were placed on thermocycler, cDNA denatured for 1 min at 98°C, followed by 1 h at 60°C to enable specific cDNA and TAC-seq probe hybridization. After hybridization, thermostable ligase reaction mixture was added. To keep a constant hybridization temperature (60°C), the cycler lid was opened and strip caps were removed. 5 µl of Taq DNA ligase master mix, containing 2× Taq DNA ligation buffer (New England Biolabs, NEB) and 1 U Taq DNA ligase (NEB) were added to each individual reaction tube and mixed by pipetting. The strip tubes were not removed from 60°C thermocycler to avoid self- and mispairing of TAC-seq probes. Ligation reaction was stopped after 20 min incubation by placing the reaction tubes on ice.

15 µl of mixture consisting of Dynabeads MyOne Carboxylic Acid beads (2 µl) (Thermo Fisher) and 13 µl of capture buffer (30% PEG-6000, 2 M NaCl, 5 mM Tris-HCl (pH 7.5), 10 mM EDTA and 0.02% Tween-20 (all chemicals from Sigma-Aldrich)) was added to ice-cooled ligated sample. The content was mixed by vortex. Capture was carried out for 10 min at room temperature. After that the tubes were placed on DynaMag-96 Side Magnet (Thermo Fisher) holding 8-well strip tubes on VersiPlate Frame (Thermo Fisher). Supernatant was removed after 3 min incubation on the magnet. The beads on magnet were washed once with 50 µl of fresh 80% ethanol. Ethanol was removed by pipetting, and the clean pellet, without ethanol drops, was dried for 2 min. Once beads were dry, strip tubes were removed from the magnet and 18 µl of PCR master mix was added directly to the beads. We have also successfully performed magnetic bead capture prior PCR without ethanol washing to avoid the risk of over-drying the bead. In the latter case, the supernatant should be removed completely. PCR master mix contained 1× proofreading HOT FIREPol Blend Master Mix (Solis BioDyne, Tartu, Estonia) and 250 nM TAC-seq left primer. In addition to universal TAC-seq left, 16 different TAC-seq barcoded oligonucleotides were used to introduce a 6-bp barcode to each studied sample (Supplemental Table S3). For that, 2 µl of 5 µM TAC-seq barcoded 1–16 primers were added individually to each PCR reaction. Strip tubes were closed with clean domed caps, mixed on vortex until beads were completely re-suspended. The ERCC spike-in reaction was incubated at 95°C for 12 min, followed by two cycles of 95°C for 20 s, 57°C for 60 s and 72°C for 20 s. In addition, 16 cycles of 95°C for 20 s, 65°C for 20 s and 72°C for 20 s with a final extension at 72°C for 1 min using the default ramp speed of the T100 cycler (Bio-Rad) were performed. PCR products were pooled together into 1.5 ml tube. The tube with pooled sample was placed on magnet to remove carboxylated beads before the following column purification. Clear supernatant was purified with DNA Clean & Concentrator-5 column (Zymo Research) and eluted with 50 µl of elution buffer (EB). The library was size-selected using AMPure XP beads (Beckman Coulter) in a single-step selection to reduce 81 bp linear PCR double-stranded by-product (Supplement Fig. S1). 50 µl beads were added to 50 µl of the purified PCR product, incubated for 5 min at room temperature and captured by a magnet for 3 min. After incubation on magnet, the supernatant was discarded and the remaining beads were centrifuged at 500×*g* for 10 s. After centrifugation, the beads were placed again on the magnet and all remaining supernatant was removed. The beads were eluted directly without ethanol washing in 25 µl of EB and incubated for 1 min at room temperature. AMPure XP bead elution has almost 100% efficiency even without previous ethanol wash. Finally, the eluted library was transferred to a clean tube after 1 min incubation on the magnet. The 180 bp library (Supplemental Fig. S1A-D) was visualized on a TapeStation High Sensitivity D1000 ScreenTape (Agilent Technologies) and quantified using KAPA Library Quantification Kit (Roche).

### Clinical sample mRNA library preparation

mRNA biomarker libraries for endometrial receptivity testing were prepared as described above with the following modifications. Total-RNA samples with RIN values 7.7–9.6 (quantified by Qubit (Invitrogen)) were diluted to concentration of 90 ng/µl and 1 µl of this was used for library preparation. RT master mix contained 1 µl of 1:50,000 of ERCC RNA Spike-In Mix 1 (Life Technologies) dilution for technical normalization. Altogether 64-plex TAC-seq probe set, containing 57 biomarker genes (Altmae et al. 2017) and seven ERCC spike-ins (ERCC-00085; 00170; 00019; 00131; 00092; 00108 and 00004) were used to generate a library for high-coverage analysis. The low-coverage analysis was performed using 70-plex, containing 57 biomarker genes (Altmae et al. 2017), five ERCC spike-ins (00131; 00108; 00092; 00019 and 00004) and eight housekeeping genes (*ACTB*, *GAPDH*, *YWHAZ*, *PPIA*, *CYC1*, *HMBS*, *TBP* and *SDHA*). 5 µM detector oligonucleotide mixtures from 100 µM stocks were created as described above. PCR was performed using in total 12 cycles, following 2+10 principle (described in details above) for both high- and low-coverage libraries.

### microRNA library preparation

miRNA profiles were analysed from endometrial total-RNA. 3’ ligation was carried out overnight in 5 µl volume. The reaction contained 100 ng of total-RNA, 1× RNA T4 RNA Ligase Reaction Buffer (NEB), 20 U RNase inhibitor (Thermo Fisher), 10% PEG-8000 (NEB), 100 nM adenylated 3’ linker and 40 U T4 RNA ligase 2 (truncated, NEB). After ligation, the free ligation adapter was removed by adding 0.5 µl 5’-Deadenylase (25 U/µl, NEB) and 0.5 µl Lambda exonuclease (5 U/µl, NEB) and incubated 10 min at 37°C, followed by 10 min at 75°C. cDNA was synthesized after adding 0.4 µl 100 mM DTT (Invitrogen), 0.4 µl 2 M KCl (Sigma-Aldrich), 0.4 µl 10 mM dNTPs (Thermo Fisher), 0.4 µl RNase inhibitor (Thermo Fisher), 0.2 µl 10 µM micro RT biotin primer and 0.2 µl Maxima H Minus Reverse Transcriptase (200 U/µl, Thermo Fisher) which were mixed into one 2 µl master mix. cDNA incubation was carried out for 15 min at 50°C, followed by 5 min at 80°C. Unbound primers were removed by adding 1 µl Exonuclease I (20 U/µl, Thermo Fisher) and incubating for 10 min at 37°C and 5 min at 95°C. 1 µl of 5 µM TAC-seq detector mixture, containing miRNA-specific left detectors and miRNA universal 5’ phosphorylated detector oligonucleotide (Supplemental Fig. S6), was added to previous 9 µl product and incubated first for 2 min at 98°C to denature the template and probes and then for 1 h at 60°C. After the hybridization, thermostable ligase reaction mixture was added on thermocycler, keeping a constant (60°C) hybridization temperature. The cycler lid was opened and strip caps were removed. 5 µl of Taq DNA ligase mixture, containing 2× Taq DNA ligation buffer (NEB) and 1 U Taq DNA ligase (NEB) was added to each individual reaction tube and mixed by pipetting. Ligation was stopped after 20 min incubation by placing reaction tubes on ice. 3 µl of Dynabeads MyOne Streptavidin C1 beads (Invitrogen) were washed according to protocol and suspended in 15 µl recommended B&W buffer. The beads were added to ligated product on ice, mixed well by pipetting and incubated for 10 min at room temperature. After capturing the beads on magnet for 1 min, the supernatant was removed and the beads were washed once with B&W buffer. TAC-seq ligated detectors were amplified as described above using 2+18 cycles of PCR. The designed miRNA library is 170 bp (Supplemental Fig. S1E).

### Cell-free DNA library preparation

10 ng of acoustically sheared (Covaris) cell-free-like genomic DNAs were combined to create excess rates of chr21 above euploid level, mimicking the extra 5–30% of fetal cfDNA fractions. 100% fraction is the GM04616 cell line’s DNA with trisomy 21. Each concentration was performed as duplicate. Samples were pipetted into strip tubes, adding 1 µl of 5 µM TAC-seq detector oligonucleotide mixture and 1 µl 10× hybridization buffer, containing 100 mM Tris-HCl (pH 7.5), 500 mM KCl, 0.2% Tween-20 and 0.1 mM EDTA. The final hybridization volume was 12 µl. The content was mixed by vortexing and centrifuged briefly. Strip tubes were placed on thermocycler, mixture denatured for 2 min at 98°C, followed by 2 h at 60°C for hybridization. After hybridization, thermostable ligase reaction master mix was added on thermocycler, keeping constant (60°C) hybridization temperature. Subsequently, 2.5 µl of Taq DNA ligase master mix, containing 1.5 µl 10× Taq DNA ligation buffer (NEB) and 1 U Taq DNA ligase (NEB) was added to each individual reaction tube and mixed by pipetting. Ligation reaction was stopped after 20 min incubation by placing reaction tubes on ice. 25 µl of previously combined Dynabeads MyOne Carboxylic Acid beads (3 µl) (Thermo Fisher) and 22 µl of capture buffer as described above was used for capture in this assay. Ligated TAC-seq detectors were amplified as above described using 2+19 PCR cycles.

### MicroRNA spike-in preparation

Custom miRNA spike-in was prepared with PCR using 76 bp synthetic ‘miRNA spike-in’ oligonucleotide, ‘TAC-seq left’ and ‘miRNA spike-in right primer’ (Supplemental Table S3). PCR was carried out in 100 µl volume containing 20 µl HOT FIREPol Blend Master Mix (Solis BioDyne), 1 µl 100 nM miRNA spike-in DNA oligonucleotide as a template, 1 µl 100 µM TAC-seq left and miRNA spike-in right primers. The reaction tube was incubated at 95°C for 12 min, followed by two cycles of 95°C for 20 s, 57°C for 60 s and 72°C for 20 s. In addition, 8 cycles of 95°C for 20 s, 65°C for 20 s and 72°C for 20 s with a final extension at 72°C for 1 min were used. The product was purified by column and quantified by KAPA Library Quantification Kit (Roche).

### Reference sequencing and data analysis

Total-RNA samples with concentration at least 200 ng/µl and RIN >8 were used for endometrial receptivity cDNA library construction. Libraries were generated from ~1 µg of total-RNA using TruSeq Stranded Total RNA (Illumina) protocol. Libraries were normalized, pooled and sequenced by Illumina HiSeq2500 instrument producing 100 cycles paired-end reads. The RNA-seq data was analyzed as previously described (Altmae et al. 2017). Heatmaps of the results were generated using the ‘pheatmap’ package implemented in R. For plotting, CPM values provided by edgeR were log-transformed, using the transformation log(CPM+1) to facilitate graphical presentation of the results.

Previously published small RNA sequencing data, containing the same RNA samples as in the miRNA TAC-seq experiments, was used. Briefly, libraries were constructed following a TruSeq Small RNA Library Preparation Guide (Illumina). 1 µg of small RNA fraction isolated from endometrial tissues was used as an input. Libraries were sequenced by Illumina HiSeq 2500 instrument producing 50 bp single-end reads. The RNA-seq data was analyzed as previously described (Altmae et al. 2017).

Sheared genomic DNA samples with concentration 5 ng/µl were used to generate cfDNA libraries as described elsewhere but using 12 cycles of PCR. Libraries were quantified by Qubit HS assay (Thermo Fisher), brought to the uniform concentration and pooled. The pooled library quality was estimated using a TapeStation High Sensitivity D1000 ScreenTape (Agilent Technologies) and sequenced by Illumina NextSeq 500 instrument producing 85 bp single-end reads. A previously described method was used for the analysis including mapping of sequencing reads to the reference genome, calculating the coverage of each region in the genome, GC correction, calculating the mean and standard deviation of the reference population and the sample. Finally, risk for aneuploidy was estimated by calculating Z-score, as well as additional ZZ-score, BM (bin median) and OM (other median). Trisomy is called if Z-score is ≥ 3, ZZ-score is ≥ 3, BM is ≥ 1.5 and OM is < 1 (Supplemental Fig. S9).

### TAC-seq sequencing

The ERCC spike-in library was sequenced by Illumina NextSeq 500 high output 75 cycles kit using 2 pM library concentration. The library was sequenced using 90-bp single-read protocol that was primed by Illumina Read1 (HP10) primer. The entire construct was 90-bp. The second, high-coverage mRNA biomarker set was sequenced with configuration identical to the one described above. In both libraries, particularly in receptivity biomarker assay, 2-channel Illumina SBS technology caused reduced level of cluster quality due to a common 20-bp motif (an extremely low-diversity region) at construct 62–82-bp site (Supplemental Fig. 8A). 4-channel SBS was used with the same library and 90-bp read using MiSeq (Illumina) instrument (data not shown) without any improvement. Following custom barcode sequencing primer was designed and used for low-coverage mRNA biomarker assay, analyzed by MiSeq Reagent Kit v3 in 14 pM library concentration. Custom barcode primer avoided the low-diversity common region and significantly improved the outcome, increasing the chastity filter (pass-filter) per cent from previous 67% to 93% (Supplemental Fig. S8B-D). In total 62-bp Read1 and 6-bp barcode (index) nucleotides were sequenced. miRNA library was sequenced by NextSeq 500 high output 75 cycles kit and 2 pM library concentration using LNA custom barcode primer. The Read1 length was 32-bp plus 6-bp barcode. Cell-free DNA library was analyzed by NextSeq500 instrument, using custom LNA barcode primer, 1.8 pM loading concentration, 62-bp for Read1 and 6-bp for barcode. The data have been deposited under GEO accession number GSE98386 and GSE110110 and SRP accession number SRP132266.

### TAC-seq data analysis

ERCC spike-in reads were trimmed to construct length of 88-bp and demultiplexed by barcodes (6-bp) allowing 1 mismatch. Demultiplexed reads were further trimmed to length of 62-bp and 4-bp of UMI at the end of the read was inserted after 4-bp of the UMI at the start of the read. Reads with UMI that contained unallocated nucleotides, were discarded. Resulting reads per sample were demultiplexed again using target regions of the genes (54-bp) allowing up to 5 mismatches. Total read counts, unique molecule counts and Pearson and Spearman correlations were calculated at different UMI thresholds. Due to the fact that Pearson method assumes linear correlation and therefore resulted in insensitive correlations at high UMI thresholds, we chose Spearman correlation as it proved to work correctly in case of very high and very low molecule concentrations.

Gene expression reads were processed as described above. To reduce potential sequencing error accumulating at UMI motif, only reads appearing at least twice were counted as unique molecules (UMI=2). Each sample was normalized to CPM using edgeR (version 3.18.1) package in R (version 3.4.1) and log-transformed using the log10 (CPM+1) transformation to reduce skewness. In addition to biomarker genes, both high- and low-coverage TAC-seq libraries included spike-in molecules. Furthermore, the low-coverage library included eight housekeeping genes which were taken into account in CPM normalization. The normalization procedure was based on the published formula that was further adjusted for read count data. Genomic DNA sequencing data quality control and pre-processing were performed as described above. Loci that were 1.5 interquartile ranges (IQRs) below the first quartile or above the third quartile were called as outliers and removed. As we constantly detected slightly higher molecule counts in chr2 compared to chr21 in euploid samples, chromosome-specific molecule counts were applied. For that, mean molecule counts of chr2 and chr21 (~1.081 at UMI threshold 2) using the euploid samples were calculated and used for normalization. User-friendly software was developed to enable TAC-seq data processing in end-user’s personal computer or in Linux environment. In both cases open-source software with installation instructions are available from https://github.com/cchtEE/TAC-seq-data-analysis and shown schematically in Supplemental Fig. S2.

### Setup and running cost of TAC-seq

Targeted assay setup needs two specific probe oligonucleotides for mRNA and cfDNA analysis. The price we got from synthesis company was 27 EUR per detector pair, resulting in 800 EUR in total for 30 loci analysis, for example (Supplemental Fig. S10A). miRNA detection needs only one non-phosphorylated specific detector per molecule of interest (Supplemental Fig. S6), lowering the setup cost to 200 EUR in case of 30 loci. Reagent costs, using off-the-shelf consumables, show that cDNA synthesis for mRNA assay is 0.6 EUR per entire sample that is more cost-effective than miRNA cDNA synthesis (3.2 EUR/sample) that needs more manipulations. Following ligation, purification and quality control cost 5.4 EUR for mRNA, miRNA and cfDNA applications (Supplemental Table S2). Sequencing contributes majority of the analysis reagent price, being in the range of 20–34 EUR per sample, depending on the application and sequencing depth (Supplemental Fig. S10B). Based on the current prices of our in-house library preparation and sequencing, the total reagent cost for miRNA profiling and cell-free DNA analysis is less than 30 EUR per sample and 26–40 EUR per sample for mRNA biomarker analysis.

### Accession codes

Supplementary sequencing data are available in Gene Expression Omnibus database under accession codes GSE98386 and GSE110110, and in Sequence Read Achieve under accession code SRP132266.

## Acknowledgements

This work was supported by the Estonian Ministry of Education and Research (grant no IUT34-16), by the Enterprise Estonia (grant no EU48695), by the EU FP7-PEOPLE-2012-IAPP, SARM (grant no EU324509), by the European Commission Horizon 2020 research and innovation programme under grant agreement 692065 (project WIDENLIFE) and MSCA-RISE-2015 project MOMENDO (grant no 691058). The authors thank S. Vuoristo for helpful discussions and M. Saare for providing information on housekeeping genes. We also thank the Karolinska Institutet Bioinformatics and Expression Analysis (BEA) core facility and High Performance Computing Centre of the University of Tartu.

## Author contributions

K.K. conceived the study.

K.K., M.K., T.J. and K.R. developed the protocols and performed TAC-seq experiments.

H.T., M.K., P.Palu., K.R. and M.P. performed TAC-seq data analysis.

T.L-P., A.V-M. and V.K. performed RNA-seq data analysis.

O.Ž. performed NIPT experiments.

J.K., A.S., and P.Palt. supervised data analysis.

H.T. and P.Palu. wrote the bioinformatics pipelines.

K.K wrote the first draft of the manuscript with support from all authors.

